# Consistent neural representation of valence in watching and recall conditions

**DOI:** 10.1101/2024.06.17.599298

**Authors:** Hyeonjung Kim, Jongwan Kim

**Affiliations:** Department of Psychology, Jeonbuk national university, Republic of Korea

**Author notes:** Corresponding author. 567 Baekje-daero, Deokjin-gu, Jeonju-si, Jeonbuk State, 54896 Republic of Korea.

**Keywords:** Stimulus presence, Recall, Regression-based decoding, Searchlight, Valence

## Abstract

Recall is an act of elicitation of emotions similar to those emotions previously experienced. Unlike the past experiences where external sensory stimuli triggered emotions, recall does not require external sensory stimuli. This difference is pertinent to the key debate in affective representation, addressing whether the representation of valence is consistent across modalities (modality-general) or dependent on modalities (modality-specific). This study aimed to verify neural representations of valence irrespective of the presence of external sensory stimuli. Using neuroimaging data from video watching and recall (Chen et al., 2017) and behavioral data for valence ratings (Kim et al., 2020), a searchlight analysis was conducted with cross-participant regression-based decoding across the presence and absence of external stimuli. Multidimensional scaling was employed as a validation analysis of the results. The searchlight analysis revealed the right middle temporal and inferior temporal gyrus as well as the left fusiform gyrus. The validation analysis further exhibited significant consistent neural representations of valence in the inferior temporal gyrus and the left fusiform gyrus. This study identified the brain regions where valence is consistently represented, regardless of the presence of external sensory stimuli. These findings contribute to debate in affective representations, by comparing conditions utilized little in prior, suggesting the inferior temporal gyrus is related to representations of valence irrespective of the presence and absence of external visual stimuli.

## 1. Introduction

### Recall and emotion

When we recall past events, we often experience emotions similar to those we experienced in the past. Tulving (2002) noted that this phenomenon occurs as we reorganize memories through the recall process. Recall has been utilized as a method of eliciting emotional experiences (Duken et al., 2021; Siedlecka & Denson, 2018), though the similarity between re-experienced emotions and past emotions varies depending on the emotions. Siedlecka and Denson (2018) reviewed various elicitation methods of being utilized in emotion studies and noted that recall for autobiographical memories triggered both positive and negative emotions (e.g., happiness, sadness) while it is challenging to elicit emotions associated with high-arousal states (e.g., surprise). These studies indicated that re-experienced emotions are similar to past emotions in that positive and negative states were similar rather than states of arousal were.

When we recall past events we experience similar emotions though, the time and situation are different. At the time of an event, we feel emotions through either the interpretation of sensory information obtained outside and the information inside the body (e.g., knowledge, fluctuation of homeostasis). In contrast, when we recall an event, no external signals or objects elicit emotions; rather we feel emotions through an integrated reconstruction of internal information obtained from past experiences (Robinson & Clore, 2002; Tulving, 2002). The idea that recall is a systematic reorganization process distinct from the information process about an experience is supported by neurological evidence. Chen et al. (2017) confirmed whether neural representations were consistent across individuals during ‘watching’ and ‘recall’ conditions. They found that neural representations during watching were consistent across individuals’ brain regions, including the low-level modality region (e.g., V1) and high-level cognition regions (e.g. the default mode network; DMN, medial prefrontal cortex; mPFC). They also found consistent neural representations across individuals who recalled and those who recalled or watched in the high-level visual cortex (e.g., the fusiform area) and high-level cognitive regions (e.g., DMN). These results indicated that there are common anatomical regions for memorizing encoded information as abstract information (Chen et al., 2017). In other words, the recall process integrally reorganizes information consistently and systematically.

### Core affect and modality

According to core affect theory (Russell, 1980), emotions are represented in two bipolar dimensions: valence, ranging from negative to positive states, and arousal, ranging from low to high states. Core affect theory is supported by behavioral, psychophysiological, and neuroimaging studies using diverse stimuli such as sounds (Bradley & Lang, 2000; Viinikainen et al., 2012), pictures (Larsen et al., 2003), music (Gomez & Danuser, 2004; Kim & Wedell, 2016), videos (Kim et al., 2017; Kim et al., 2020), and recall (Gomes et al., 2013; Kensinger & Corkin, 2004).

According to Barrett and Bliss-Moreau (2009), the neural network of core affect is a primary component of neural circuits integrating external sensory information with information from inside the body. The neural networks of core affect can be subdivided into two functional networks (Barrett & Bliss-Moreau, 2009). The first is a sensory integration network connecting unimodal brain areas related to sensory modalities. This network processes object representation such as visual features, together with the effect on the body’s homeostasis. The second is a visceromotor network that is involved in the behavioral, hormonal, and autonomic responses to the object. Based on these two different functional networks, two hypotheses were proposed for affective representations across modalities (Kim et al., 2017). The first hypothesis is modality-specific, corresponding to the sensory integration network, and argues that affective representations are unique for each modality. The second hypothesis is modality-general, corresponding to the visceromotor network, and claims that affective representations are consistent (universal) across modalities.

Whether valence is represented consistently or uniquely across modalities has been debated because neurological evidence has been reported for both hypotheses. Within the core affect theory, valence representations are thought to be modality-general (Kober et al., 2008), and various studies support this (Chikazoe et al., 2014; Dalenberg et al., 2018; Kim et al., 2017). However, there is also evidence for the modality-specific hypothesis (Miskovic & Anderson, 2018; Shinkareva et al., 2014), plus results supporting both hypotheses have been reported (Gao & Shinkareva, 2021).

### Recall within the core affect theory

The relationship between emotions elicited by experience and recall can be explained by the core affect theory and affective representation hypotheses for modality. The similarity between emotional states triggered by experience and recall suggests similar representations on the valence dimension of core affect. The different processes of eliciting emotion are related to the affective representation hypotheses for modality. When valence is elicited by experience, external stimuli are present. Valence may be uniquely or consistently represented depending on the stimuli’s sensory modalities. When valence is elicited by recall, there are no external stimuli or specific sensory modalities present. Thus, valence can be represented consistently regardless of external modalities. The emotions elicited by experience and recall are felt similarly states suggests that valence is represented consistently regardless of external sensory stimuli. This assertion is indirectly supported by a study (Duken et al., 2021), which observed consistent physiological emotional responses regardless of the presence of external sensory stimuli.

Prior studies typically compared valence representations across different modality stimuli (e.g. visual and auditory). Research comparing recalled and past-experienced valence investigated the presence or absence of external modality rather than different modalities. For example, Skerry and Saxe (2014) manipulated the presence of direct evidence for understanding other people to find common neural representations of perceived and inferred emotion. They found consistent neural representations for inferred and perceived emotion with and without direct stimuli.

Our study aimed to verify whether neural representations of valence are consistent or specific according to the presence or absence of external stimuli. We used watching and recall conditions to compare the valence elicited from an external modality with the valence elicited from internal, integrated information. There is limited evidence from neuroimaging on whether the valence representations during recalling and watching involve a common neural code. Therefore, we used neuroimaging data to explore brain regions that showed consistent neural representations of valence elicited during watching and recalling. Whereas watching audiovisual videos elicits emotion through an external sensory experience, recall elicits re-experienced emotion through an integrated reorganization of internal sensory and abstract information, without external stimuli. Consistent patterns of neural representations of valence during watching and recall across the brain regions would support the assertion that valence is represented consistently regardless of the presence of external stimuli.

### Naturalistic stimuli

Experimental stimuli used to elicit emotions in previous studies primarily focused on excluding the influence of control variables. The stimuli typically comprised single images, videos several seconds in length, or sounds designed in advance to suit the researchers’ purposes. Experimental stimuli increase an experiment’s internal validity but reduce its external validity because of discrepancies between the experimental and real-world environments. To overcome this, emotion studies have recently started using naturalistic stimuli. Naturalistic stimuli resemble events one might encounter in real life, including various sounds, scenes, and context. Because naturalistic stimuli have the advantage of similarity to experiences in the actual living environment (Schmuckler, 2001), they are utilized in cognitive neuroscience and emotion studies to improve ecological validity (Chang et al., 2021; Hasson et al., 2010; Hasson et a., 2008). As recall is the process of retrieving and reorganizing various information stored from past experiences (Robinson & Clore, 2002; Tulving, 2002), naturalistic stimuli are suitable for investigating recall.

### Multivariate pattern analysis

We performed multivariate pattern analysis (MVPA) due to its advantages over univariate analysis in fMRI studies. While univariate analysis is useful in confirming the interaction effects and differences between groups, it does not consider the covariates between dependent variables. This analysis potentially increases the error level of a study’s results when comparing multiple groups or conditions repeatedly, called the multiple comparisons problem. Unlike univariate analysis, MVPA is suitable for analyzing numerous data, is not affected by the multiple comparisons problem, and minimizes the information lost through excessive summarization.

One MVPA method, decoding analysis, uses data patterns to predict categories and values and has been utilized in emotion studies examining whether neural patterns explain affective representations (Baucom et al., 2012; Weaverdyck et al., 2020). There are two types of decoding analysis: regression-based decoding and classifications based on machine learning (Weaverdyck et al., 2020). Classification, which identifies categorical conditions or variables, rather than continuous variables, has been used in emotion studies using behavioral (Kim et al., 2020), psychophysiological (Mower et al., 2011; Verma & Tiwary, 2014), and neuroimaging data (Baucom et al., 2012; Li et al., 2019; Skerry & Saxe, 2014). To apply classification, the stimuli must be assigned to each condition decided by the researcher in advance. In contrast, regression-based decoding enables predictions of continuous variables, such as participants’ ratings (Kim et al., 2020), and the stimuli do not require assigning to each condition. We applied regression-based decoding to predict the participants’ valence ratings based on neural representations.

Searchlight analysis was used to explore the anatomical regions exhibiting valence representations consistently across participants in the presence and absence of external stimuli. Searchlight analysis produces a whole-brain map consisting of particular values (e.g., correlation values) (Kriegeskorte et al., 2006). Statistical analysis is conducted using neighboring voxels located within a specified area centered on each voxel. By assigning obtained statistics to a center voxel, the brain map is constructed. Searchlight analysis has been used to explore specific regions of interest in the brain (Kim et al., 2017; Visconti di Oleggio Castello et al., 2021) and boasts the advantages of MVPA while overcoming the disadvantage that MVPA does not reveal spatial information about neural activity. We performed searchlight analysis with regression-based decoding to produce brain maps showing the prediction accuracy (correlation values) of participants’ valence. Additionally, we conducted the permutation test to whether the clusters identified by the searchlight analysis were significant. The permutation test is used to check for significance because it produces the null distribution by shuffling the data randomly when there is no known null distribution for specific statistics. This method enables researchers to test the statistical significance of experimental results, even when using nonparametric testing methods or designs where bias can occur.

Multidimensional scaling (MDS) was conducted to validate the significant clusters identified from the searchlight analysis. In MDS, each object is represented in the low-dimensional space, utilizing dissimilarity data about high-dimensional stimuli (Shinkareva et al., 2013). Then, we employed Procrustes rotation to place the stimuli in the low-dimensional space based on valence hypotheses, and based on the presence or absence of external stimuli: (1) the presence–absence dimension, which defined the watching and recall conditions; (2) the presence–absence general dimension, which hypothesizes consistent valence representations across two conditions; and (3) the presence–absence specific dimension, which hypothesizes distinctive valence representations across the two conditions. If brain regions found to be significant exhibited consistent valence representations regardless of the presence of external stimuli, then neural representations in those regions were expected to discriminate between emotional stimuli in a low-dimensional space based on the presence–absence general dimension while failing to discriminate between emotional stimuli in a low-dimensional space based on the presence–absence specific dimension.

### 2. Method

This study reanalyzed fMRI data from Chen et al. (2017) and behavioral ratings for affective representations from Kim et al (2020).

### Prior studies’ experiments

Chen et al. (2017) recruited 22 participants (12 male, 10 female, aged 18–26 years, mean age: 20.8 years) who had not watched the first episode of the British television series *Sherlock*. Inside the fMRI scanner, participants watched the first *Sherlock* episode, divided into two videos 23 and 25 minutes long, respectively. Before watching each of two videos, participants watched the cartoon (*Let’s All Go to the Lobby*) unrelated to the *Sherlock*, as the onset of stimulus may increase participants’ arousal states generally. After watching the videos, participants were asked to verbally recall what they remembered about the episode in narrative order and as much detail as possible for at least 10 minutes. They were encouraged to mention any additional moments they remembered out of sequence and were asked to indicate when they were finished. Participants’ fMRI data were measured while they watched and recalled the videos (Fig. 1A). Five participants’ data were excluded from the dataset because of sleep or head movement.

**Fig. 1.**
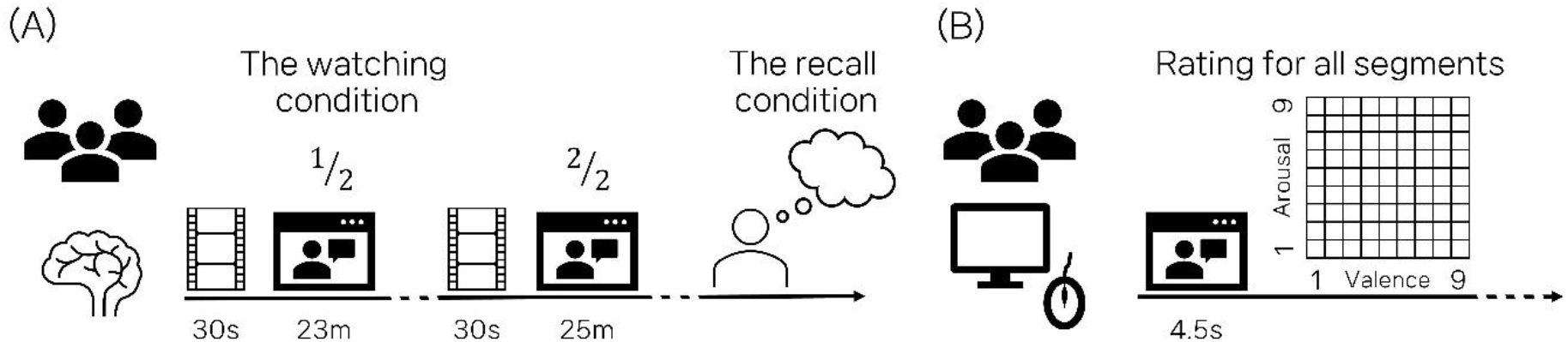
Illustrations of (A) Chen et al.’s (2017) and (B) Kim et al.’s (2020) experimental procedures.

Kim et al. (2020) also utilized the first episode of *Sherlock*, dividing the program into 621 of 4.5-s segments. Participants rated their valence and arousal state as they watched each segment using a mouse on a 9×9 two-dimensional grid. The horizontal axis represented valence, ranging from 1 (negative) to 9 (positive), and the vertical axis represented arousal, ranging from 1 (low arousal) to 9 (high arousal) (Fig. 1B). Each video segment was rated by a total of 125 participants.

### fMRI data acquisition and preprocessing

Chen et al. (2017) acquired all MRI data on a 3T Siemens Skyra scanner with a 20-Channel head coil. Functional images were collected using a T2-weighted echo-planar imaging (EPI) pulse sequence: TR = 1,500 ms, TE = 28 ms, flip angle = 64, whole-brain coverage = 27 4-mm thick slices, in-plane resolution = 3×3 mm^2^, FOV 192×192 mm^2^, with ascending interleaved acquisition. Whole-brain anatomical images were obtained using a T1-weighted magnetization-prepared rapid gradient echo (MPRAGE) pulse sequence (size of voxel = 0.89 mm^3^ resolution).

Chen et al.’s (2017) functional images were preprocessed using FMRIB Software Library (FSL). Preprocessing included slice-timing difference correction, motion correction, linear detrending, high-pass filtering (140s cut-off), and coregistration and affine transformation of functional volumes to a template (Montreal Neurological Institute standard 152). Functional images were resampled to 3 mm isotropic voxels for all analyses. Preprocessed data were standardized across time for every voxel and then smoothed with a Gaussian kernel of 6 mm full width at half maximum (FWHM). Head motion when measuring the fMRI data was minimized by instructing the participants to stay still and using foam padding in the recall session.

### Data reorganization

Chen et al. (2017) analyzed fMRI data collected while watching the 48 scenes of the Sherlock videos used in the experiment and two scenes from the two cartoons—50 scenes in total, separated by narrative. The fMRI data for each scene was obtained by averaging the brain imaging data measured during each scene. We used the fMRI data for the 48 *Sherlock* scenes, excluding the cartoons. We also excluded fMRI data measured during watching for a scene for each participant, if that scene was not successfully recalled. All volumes were standardized across scenes.

Kim et al. (2020) divided the *Sherlock* video into 621 segments, and participants rated their valence and arousal for each segment. We utilized these behavioral ratings for valence to compute the valence values for the 48 scenes as divided by Chen et al. (2017). In our study, the valence values for each scene were calculated by averaging the behavioral ratings for valence for the corresponding segments. To reconstruct a scale ranging from −4 to 4, we subtracted the obtained valence values from 5, which represents the median of the scale used in Kim et al. (2020). Negative values indicated negative valence and positive values indicated positive valence. For Chen et al.’s (2017) each participant, we rearranged the scenes according to their neural activation data from the most negative to the most positive scene. This reordering process was applied to all participants and conditions (watching and recall). Subsequent analyses were then performed using the reordered data.

### Searchlight analysis

To explore the brain regions consistently exhibiting neural representations of valence irrespective of external stimuli, we performed searchlight analysis with cross-participant cross-condition regression-based decoding (Fig. 2). First, the participants from Chen et al. (2017) were arranged into two groups: a training group and a testing group. Of 30 participants, 29 were in the training group, leaving one in the testing group. Data from a participant in each group were extracted from the voxels located within a cubic 5×5×5 volume (voxel-unit) centered on one voxel at the same coordinate across participants (Fig. 2A). The voxels’ activations measured while watching the videos and during recall were extracted for each group. To implement a cross-condition (watching and recall) paradigm, the testing group’s voxel data measured while watching were used for analysis when the training group’s voxel data during recall were used for analysis.

**Fig. 2.**
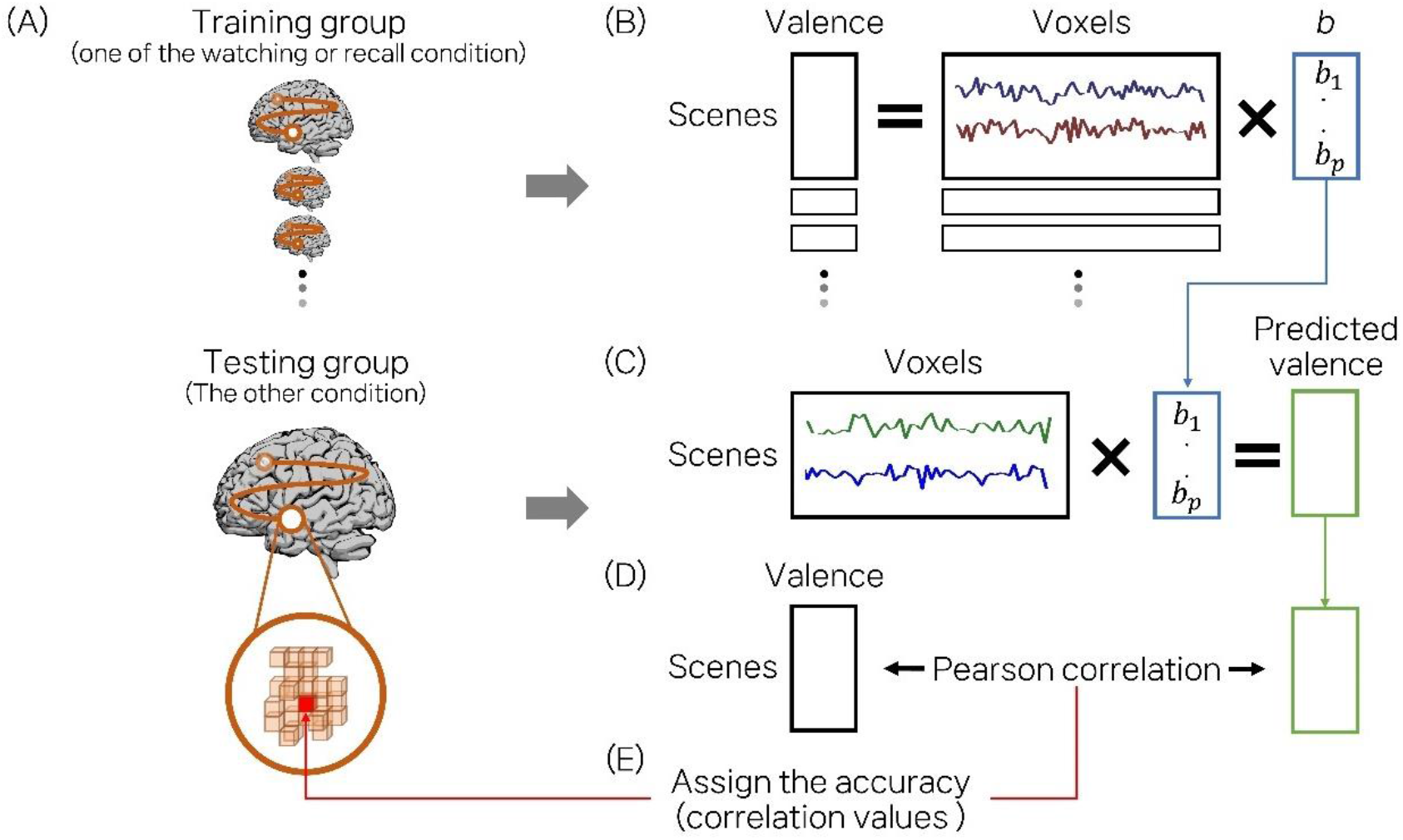
Searchlight analysis process.

Next, regression-based decoding was performed by using the training group’s voxel data as the independent variable and valence values for the scenes (as the dependent variable (Fig. 2B). By multiplying the regression coefficients for the voxels and the testing group’s voxel data, we calculated the predicted values for the testing group’s valence (Fig. 2C). Then, the Pearson correlation coefficient was calculated to obtain the degree of consistency between the predicted valence values and the testing group’s actual valence values for each scene they successfully recalled (Fig. 2D). We considered correlation values as prediction accuracy because these indicate similarity between predicted values and actual values. We arranged the correlation values in the testing group’s center voxel (Fig. 2E). We conducted this procedure for the whole brain to produce the testing group’s brain map comprising prediction accuracies (correlation values). This procedure was performed once for each condition for a participant in the testing group. Because there were data from two conditions, we obtained two maps for a participant. We averaged the two brain maps to produce an individual’s brain map, which consisted of average prediction accuracies of different prediction directions. This procedure was repeated for all participants until each of them were arranged in the testing group once.

All the brain maps were submitted for group analysis using Statistical Parametric Mapping 12 (SPM12). As per previous studies (Kim et al., 2017; Woo et al., 2014), We performed the one-sample t-test (uncorrected α = .001) and explored significant clusters (cluster size > 40).

To test the significance of the searchlight analysis’s results, the permutation test was performed. We randomly shuffled the data (the training group participants’ neural activation and scene valence ratings) to randomize the relationship between neural activation and valence levels for each participant. We did the same for the test group’s data. Then, we performed searchlight analysis using the same procedure on the shuffled dataset. We recorded the largest cluster size and repeated the entire procedure 1,000 times. We yielded a null distribution for cluster size by using 1,000 values for cluster size. The critical value for the significant cluster size was in the ninety-fifth percentile of the null distribution (α = .05).

To report the anatomical areas involved in each cluster, we reported the clusters as anatomical regions, including the peak voxel, based on the automated anatomical labelling atlas 3 (AAL3) (Rolls et al., 2020). Where the number of voxels belonging to the same region as the peak voxel did not satisfy our study’s pre-set cluster size condition (cluster size > 40), we reported the region that consisted of more than 40 significant voxels together with the region the peak voxel belonged to.

### Multidimensional scaling

Multidimensional scaling as validation analysis was conducted where the searchlight analysis identified a significant cluster. In the searchlight analysis, valence ratings for scenes that one individual successfully recalled were predicted by utilizing a regression model fitted with the other participants’ neural representations in the other condition. Therefore, a significant cluster implied consistent neural representations of valence across participants and a general pattern across condition irrespective of external stimuli. Due to variations in the number of scenes successfully recalled by each participant, the number of scenes for which valence was predicted varied across individuals. Hence, three-dimensional multidimensional scaling (MDS) was conducted to validate whether the clusters found by the searchlight analysis were general neural representations of valence irrespective of external stimuli (Fig. 3). The voxel data for each scene was averaged across all participants because not all participants recalled every scene. This process was conducted separately for the watching and recall conditions. Subsequently, correlation analysis between the 48 scenes (watching; presence condition) and 48 scenes (recall; absence condition) was performed by using voxel data from each scene measured under the two conditions. This resulted in a correlation matrix consisting of 96×96 (48 scenes watched and 48 scenes recalled). This correlation matrix (similarity matrix for stimuli) was transformed into a dissimilarity matrix for stimuli by subtracting it from one. The dissimilarity matrix was then submitted to MDS.

**Fig. 3.**
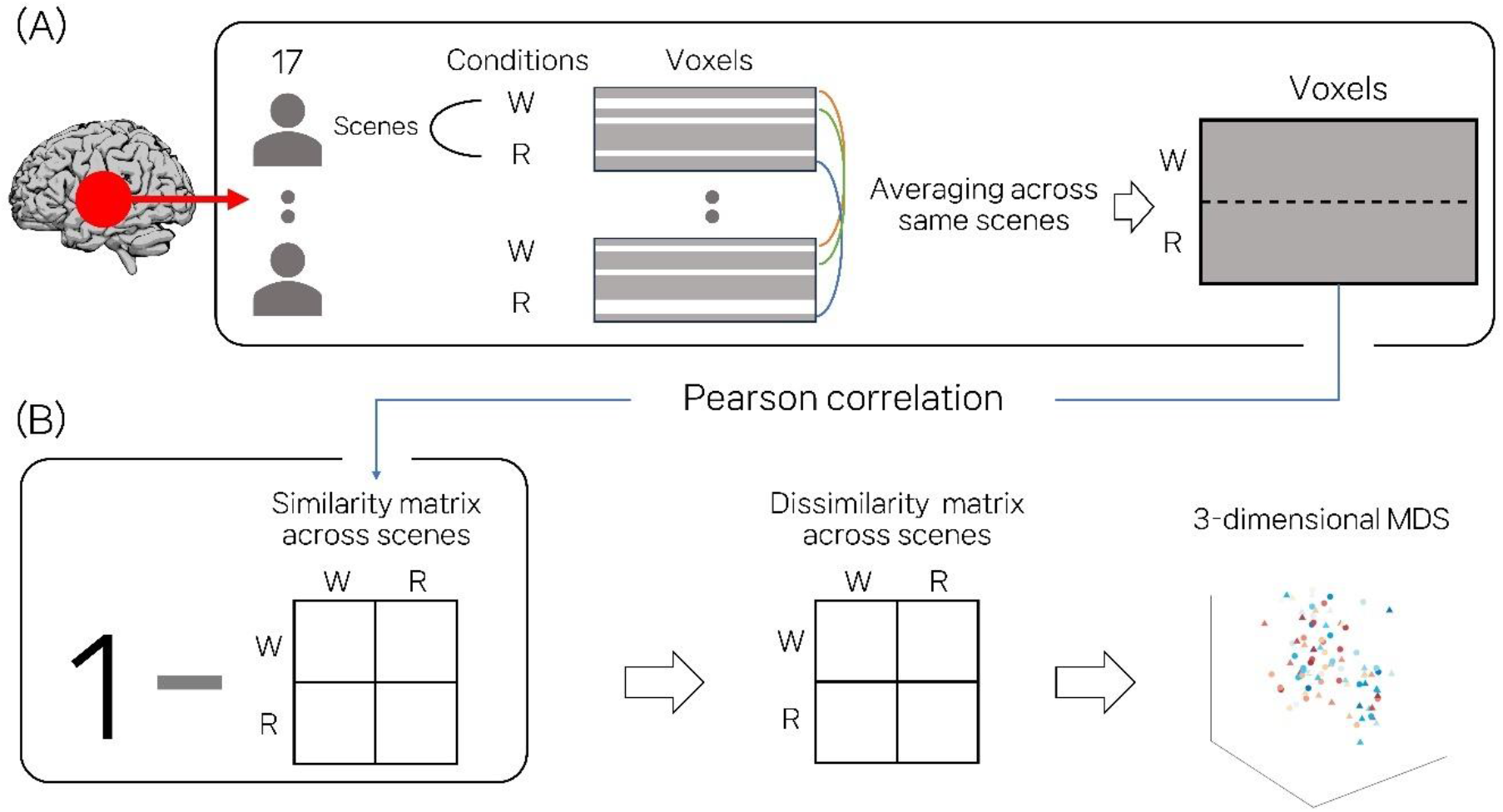
Multidimensional scaling process, where ‘W’ indicates the watching condition and ‘R’ indicates ‘recall’ condition. The colored line connects the same scenes under different conditions.

After MDS, we conducted Procrustes rotation to represent each scene’s three-dimensional solution coordinates obtained from the three-dimensional MDS in the presence–absence, presence–absence general, presence–absence specific dimensions. We used design values for presence–absence and target coordinate values based on the hypotheses for presence–absence general and specific (Fig. 4). All 96 scenes were represented in one two-dimensional space consisting of the presence–absence×presence– absence general dimensions and another consisting of the presence–absence×presence–absence specific dimensions. Design values and target coordinate values, pre-set for Procrustes rotation, were prepared for all three dimensions. First, the design values for presence–absence were constructed to verify whether neural representations differentiated between watched and recalled scene. Recalled scenes were assigned a design value of –1, and watched scenes were assigned a design value of 1. Second, because the presence–absence general dimension assumed consistent representations of valence irrespective of external stimuli, valence ratings obtained for each scene, regardless of the conditions, were set as target coordinate values. For contrast, valence ratings were z-scored across scenes. Lastly, the presence– absence specific dimension assumed distinctive representations of valence based on the presence of external stimuli. Therefore, valence ratings with opposite signs between presence and absence, were set as target coordinate values, by multiplying design values for the presence–absence condition and target coordinate values for presence–absence general. To assess the separation of scenes in each dimension based on these hypotheses, correlation analyses were conducted between the three-dimensional solution coordinates for each scene obtained through the Procrustes rotation and the pre-set design and target coordinate values for each dimension. This process was consistently applied to all the significant clusters identified by the searchlight analysis. Through this, we aimed to find clusters that significantly differentiated scenes in the presence-absence general dimension but not in the presence-absence specific dimension, confirming the corresponding brain regions.

**Fig. 4.**
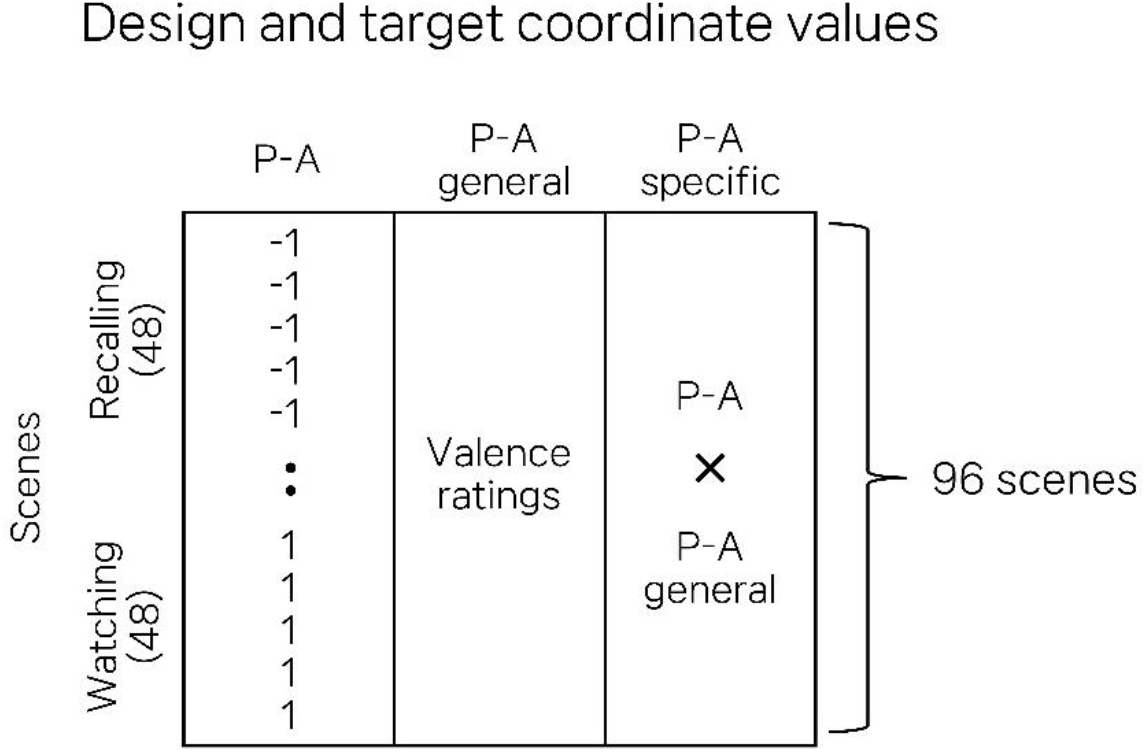
Design and target coordinates matrix. P-A means presence–absence, indicating the presence and absence of external stimulation.

## 3. Results

### Data reorganization

We reorganized the Chen et al.’s (2017) fMRI data by the number of scenes each participant successfully recalled (*M* = 34, range = 24 ∼ 46). We also restructured the valence ratings of the scenes successfully recalled by each participant, to determine if recall was biased toward positive or negative scenes. All participants successfully recalled scenes that included negative, neutral, and positive valence (range of average valence rating = –2.98 to 1.08) (Fig. 5).

**Fig. 5.**
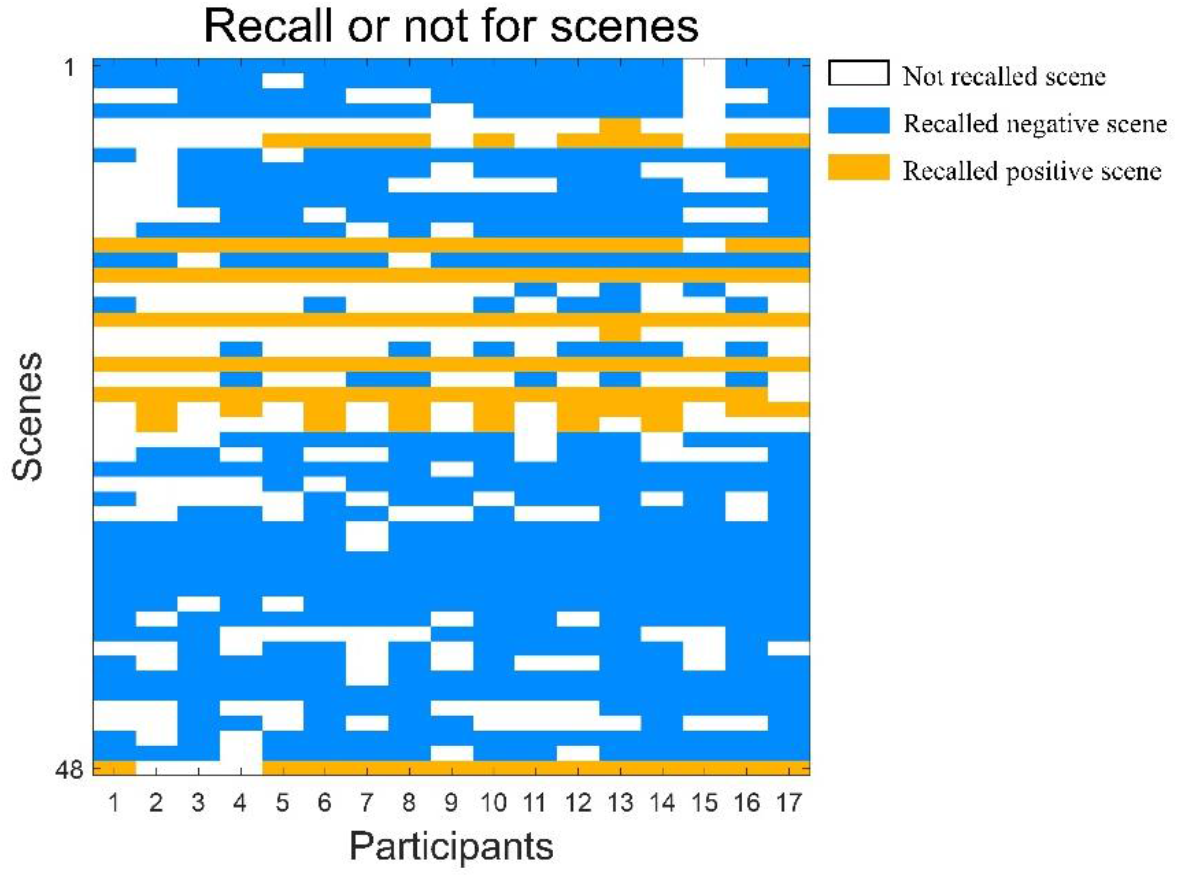
Diagram detailing the participants’ recall success. The x-axis represents individual participants, while the y-axis corresponds to each scene.

### Searchlight analysis

We conducted searchlight analysis with regression-based decoding and permutation to explore the brain regions revealing valence representations consistently across participants, irrespective of the presence of external stimuli. Three brain regions were identified: the right middle temporal gyrus (MTG), the right inferior temporal gyrus (ITG), and the left fusiform gyrus/inferior temporal gyrus (*ps* < .05, cluster size > 174, Table 1, Fig. 6, Supplementary Table 1).

**Table 1.** Significant clusters (*ps* < .05) from searchlight analysis.

| Region | Hemisphere | Cluster size | Peak voxel coordinate (MNI 152) | | | $t$ |
| --- | --- | --- | --- | --- | --- | --- |
|  |  |  | X (mm) | Y (mm) | Z (mm) |  |
| Middle temporal gyrus | Right | 180 | 63 | -15 | -24 | 7.00 |
| Inferior temporal gyrus | Right | 194 | 63 | -57 | -9 | 6.47 |
| Fusiform gyrus | Left | 178 | -27 | -3 | -39 | 6.40 |
| Inferior temporal gyrus |  |  |  |  |  |  |

Each cluster was labeled as a region based on AAL3. Peak voxel coordinate was followed by MNI 152 template. Statistic ( $t$ ), which means the peak $t$ value in the cluster, was obtained from one-sample $t$ -test using SPM12.

**Fig. 6.**
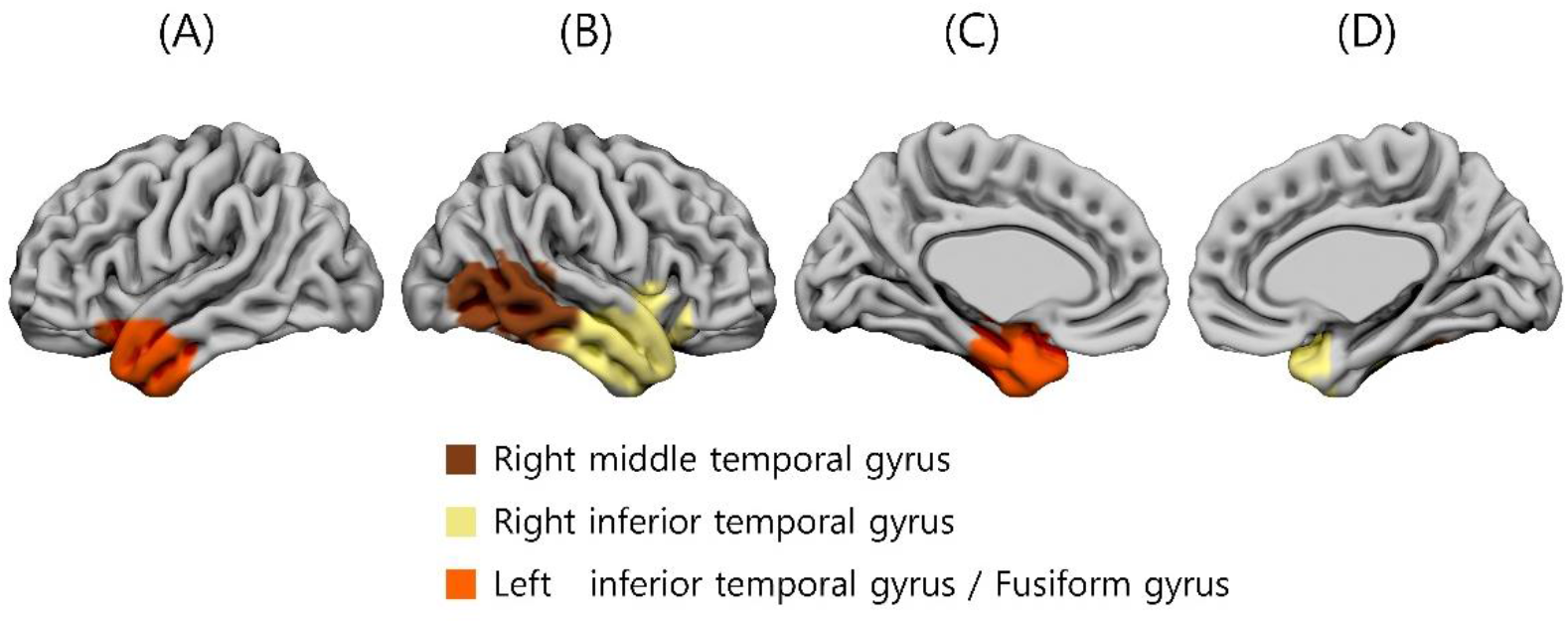
Brain regions identified by the searchlight analysis as having significant clusters. (A) and (C) represent the left hemisphere, (B) and (D) correspond to the right hemisphere. (A) and (B) represent the lateral view. (C) and (D) represent the medial view.

### Multidimensional scaling validation analysis

To verify whether the clusters found through the searchlight analysis were related to consistent valence representations across participants as well as regardless of external stimuli, we performed a three-dimensional MDS and Procrustes rotation on each cluster. We also performed a correlation analysis between the three-dimensional MDS solution values for each scene obtained after the Procrustes rotation and values of design and target coordinate for each dimension (Table 2). First, the right middle temporal gyrus (MTG) did not significantly differentiate scenes on all dimensions (*ps* > .05). Second, the right inferior temporal gyrus (ITG) differentiated scenes significantly on the presence–absence general dimension (*r* = .23, *p* < .05) but not on the presence–absence and presence– absence specific dimensions (*ps* > .05). Finally, the left fusiform gyrus/inferior temporal gyrus differentiated scenes significantly on the presence–absence general (*r* = .24, *p* < . 05) and presence– absence dimensions (*r* = .27, *p* < .01) but not the presence–absence specific except dimension (*p* > .05).

**Table 2.** Pearson correlation values between MDS solution values and either design or target coordinate values for scenes in each dimension.

| Region | Hemisphere | 3 dimensions |  |  |
| --- | --- | --- | --- | --- |
|  |  | P–A | P–A general | P–A specific |
| Middle temporal gyrus | Right | .16 | .05 | .10 |
| Inferior temporal gyrus | Right | .05 | .23* | .06 |
| Fusiform gyrus | Left | .27** | .24* | .09 |
| Inferior temporal gyrus |  |  |  |  |

\* $p < .05$ , \*\* $p < .01$ . P–A means presence and absence of external stimulation.

## 4. Conclusion

This study explored the evidence for consistent neural representations of valence regardless of the presence or absence of external modality. The results show that the right middle temporal gyrus (MTG), right inferior temporal gyrus (ITG), and left fusiform gyrus/inferior temporal gyrus are involved in consistent neural representation of valence across participants irrespective of external stimuli. Additionally, the the validation analysis using MDS revealed that the right ITG and left fusiform gyrus/ITG did significantly explain general valence representations.

### Significant brain regions in both analyses

The bilateral ITG and the left fusiform gyrus showed significantly consistent neural representations of valence across presence and absence of external stimuli in both the searchlight and validation analysis. The ITG is involved in the representation of negative and positive valence (Blair et al., 2007; Goldin et al., 2008) and various emotional functions, including emotional expression (Deng et al., 2013; Lin et al., 2020), facial emotion recognition (Dominguez-Borras et al., 2009; Morningstar et al., 2020), and emotion regulation (Lin et al., 2020). During recognition and recall, the ITG receives information from the hippocampus and prefrontal cortex (Langleben et al., 2009; Tomita et al., 1999). This region is known to play an important role in maintenance, storage, and visual representations during visual recognition and recall (Axmacher et al., 2008; Nakamura et al., 2000; Petersson et al., 1999; Ranganath et al., 2004).

In this study, bilateral ITG was activated and these activations predominantly located in the anterior temporal lobe. The anterior temporal lobe is visual and auditory integration hub and is reportedly involved in semantic memory and processing (Visser & Lambon Ralph, 2011). Visser and Lambon Ralph (2011) examined the function of the anterior temporal lobe at a subregional level and asserted that the inferior anterior temporal gyrus is involved in the integrated semantic processing of multimodalities. They further claimed that the processing of auditory information in the inferior temporal gyrus is influenced by other regions involved in auditory-information processing (e.g., the superior anterior temporal gyrus). The ITG is particularly involved in visual processing (Jomori et al., 2013; Lin et al., 2020), and its ability to process visual stimuli is modulated by changes in the auditory environment or emotion evoked by auditory elements (Dominguez-Borras et al., 2009; Jomori et al., 2013). In short, the ITG is a specifically involved in visual processing, and this is why it is thought to be involved in valence representation, recognition, and recall. During recall, emotional re-experience occurs as a result of reorganizing past information (Tulving, 2002). Thus, consistent neural representation of valence in the ITG, regardless of the presence of external stimuli, are due to the processing of visual information stored while watching.

The fusiform gyrus is related to the visual cortex and limbic areas (Jomori et al., 2013), particularly regarding in the recognition of facial expressions (Dominguez-Borras et al., 2009; Sato et al., 2004). The fusiform gyrus is sensitive to dynamic facial expression changes (Sato et al., 2004). Our study used data obtained from the naturalistic stimuli featuring actors’ faces, which might have activated the fusiform gyrus. Each scene is distinguished well by the left fusiform gyrus/inferior temporal gyrus clusters in both the presence–absence general dimensions and the presence–absence dimensions. This could be due to the absence of external visual stimuli to recognize in during recall in contrast to when watching. The finding that a cluster including the left fusiform gyrus/inferior temporal gyrus discriminated between the presence and absence of external stimuli is likely due to the fusiform gyrus’s association with visual features (e.g., face).

### Significant brain region in only searchlight analysis

Searchlight analysis identified the right middle temporal gyrus as involved in consistent neural representations of valence, but this finding was not validated. The right middle temporal gyrus is related to emotion (Dalenberg et al., 2018; Gao & Shinkareva, 2021; Lindquist et al., 2016), language (McDermott et al., 2003), memory (Wei et al., 2012), and social interaction (Sato et al., 2012). There have been trials subdividing (e.g. lateralizing) the middle temporal gyrus’s functions because it has so many. For example, language processing was lateralized in the left middle temporal gyrus (Xu et al., 2019; Xu et al., 2015). This brain region is involved in visual properties and relational categories processing (Leshinskaya & Thompson-Schill, 2020). In particular, Leshinskaya and Thompson (2020) reported that representations for visual information emerged during visual processing of events were involve in anterior to posterior regions of the right middle temporal gyrus, representing generalizable and specific visual relations. We found significant neural activations related to consistent valence representations irrespective of external stimuli and even across participants, but we did not find significantly consistent neural representation for valence in the validation analysis. There is no the clear evidence indicating that the right MTG might be involved in processing visual information and valence representations regardless of external sensory stimuli. Further studies using various modalities (e.g. visual, auditory, taste, etc.) are needed to confirm valence representation within the middle temporal gyrus.

### Modality-general hypothesis

A debate within the affective neuroscience pertains to whether the valence states are represented consistently or specifically depending on the external stimuli’s modalities (Barrett & Bliss-Moreau, 2009). The modality-general hypothesis asserts that valence states are represented consistently regardless of the modalities eliciting the emotions, while the modality-specific hypothesis claims that valence states are represented specifically depending on the modalities. To answer this debate, patterns of neural representations of valence elicited by different modalities (e.g. visual and auditory) have been explored (Gao & Shinkareva 2021; Kim et al., 2017). This study confirmed consistent neural representations of valence regardless of the presence or absence of external modalities. For comparison, we used neural representations during watching and recall and found that the inferior temporal gyrus is involved in the consistent neural representation of valence regardless of the presence of external stimuli.

Valence was elicited while watching videos, through the experience of external sensory stimuli. In contrast, recall elicits valence through the integration and reconstruction of internal and abstract information without external sensory experiences. The results indicated that the neural representations of valence in the inferior temporal gyrus occur consistently regardless of the presence of external sensory stimuli, suggesting that the inferior temporal gyrus represents emotions based on the presence– absence general dimension. The confirmation that the inferior temporal gyrus represents valence consistently, supporting the modality-general hypothesis.

Our study complements a limitation of previous research. Skerry and Saxe (2014) identified the activation of regions of interest for emotion recognition and inference using videos of facial expressions, animated videos depicting social situations, and positive or negative rewards. They found that neural activation in regions of interest was consistent regardless of the presence of direct cues for emotion recognition. However, there is little evidence that was found for consistent activation patterns outside of the region of interest when they performed the whole-brain searchlight analysis. A recent study (Duken et al., 2021) compared valence representations during recall and watching using physiological data, but neurological studies were needed. Our study contributes to the further exploration of other regions that show consistent neural representations of valence regardless of the presence of external stimuli, and provides neural evidence for consistent representations across watching and recall conditions.

### Limitation

First, our study is limited by its use of data obtained from naturalistic stimuli. Naturalistic stimuli have advantages such as including scenes with sounds and narrative contexts developing over time (Schmuckler, 2021). However, it is difficult for researchers to control the conditions eliciting the participants’ emotions using these stimuli. In our study, negative scenes were more successfully recalled by participants than positive and neutral scenes (Fig. 5).

Second, although our naturalistic stimuli were both visual and auditory, our results primarily showed activation of brain regions involved in visual processing. One assumption is that this is due to the relationship between recall and imagery. Auditory memory is inferior to visual memory (Cohen et al., 2009), and how well people imagine affects their ability to recall. (Marks, 1973; Penney, 1989). We only used data from scenes recalled successfully, meaning presence-absence of external modality is less likely to be found in brain regions involved in auditory features than in those involved in visual features. The other assumption is that the brain regions related to visual processing were mainly activated by the visual dominance. When stimuli consisted of both auditory and visual components, people refer to the visual rather than the auditory information (Colavita, 1974). Furthermore, Posner et al. (1976) demonstrated that when people pay attention to visual information to process it, they give less attention to information from the other modalities. Naturalistic stimuli have various visual scenes including facial expressions and places.

Plus, we only used data from successfully recalled scenes for the searchlight analysis. Thus, it is possible that visual processing was more dominant than auditory processing, resulting in identification of the brain regions involved in visual processing. For this limitation, we suggested future studies considering designs that equally control for the visual component and manipulate the presence of the auditory component.

