## supplementary Table 1 for "Consistent neural representation of valence in watching and recall conditions"

**Supplementary Table 1.** The anatomical regions' labels for voxels of each cluster based on Automated anatomical labeling atlas 3 (AAL3).

| | Region | The number of voxels | | Region | The number of voxels |
|---|---|---|---|---|---|
| Cluster 1 | Temporal_Mid_R | 57 | Cluster 3 | Temporal_Inf_L | 71 |
| | Temporal_Pole_Sup_R | 40 | | Temporal_Pole_Mid_L | 30 |
| | Tempporal_Pole_Mid_R | 40 | | Temporal_Pole_Sup_L | 21 |
| | Temporal_Inf_R | 30 | | ParaHippocampal_L | 17 |
| | OFCpost_R | 6 | | Fusiform_L | 12 |
| | Insula_R | 5 | | Temporal_Mid_L | 12 |
| | Frontal_Inf_Orb_2_R | 2 | | Amygdala_L | 7 |
| Cluster 2 | Temporal_Mid_R | 106 | | Hippocampus_L | 3 |
| | Temporal_Inf_R | 85 | | Insula_L | 2 |
| | Occipita_Inf_R | 2 | | OFCpost_L | 1 |
| | Fusiform_R | 1 | | | |
